# Bioactivity of curcumin on the cytochrome P450 enzymes of the steroidogenic pathway

**DOI:** 10.1101/669440

**Authors:** Patricia Rodríguez Castaño, Shaheena Parween, Amit V Pandey

## Abstract

Turmeric, a popular ingredient in the cuisine of many Asian countries, comes from the roots of the *Curcuma longa* and is known for its use in Chinese and Ayurvedic medicine. Turmeric is rich in curcuminoids, including curcumin, demethoxycurcumin, and bisdemethoxycurcumin. Curcuminoids have potent wound healing, anti-inflammatory, and anti-carcinogenic activities. While curcuminoids have been studied for many years, not much is known about their effects on steroid metabolism. Since many anti-cancer drugs target enzymes from the steroidogenic pathway, we tested the effect of curcuminoids on cytochrome P450 CYP17A1, CYP21A2, and CYP19A1 enzyme activities. When using 10 µg/ml of curcuminoids, both the 17α-hydroxylase as well as 17,20 lyase activities of CYP17A1 were reduced significantly. On the other hand, only a mild reduction in CYP21A2 activity was observed. Furthermore, CYP19A1 activity was also reduced up to ~20% of control when using 1-100 µg/ml of curcuminoids in a dose-dependent manner. Molecular docking studies confirmed that curcumin could dock into the active sites of CYP17A1, CYP19A1 as well as CYP21A2. In CYP17A1 and CYP19A1, curcumin docked within 2.5 Å of central heme while in CYP21A2 the distance from heme was 3.4 Å, which is still in the same range or lower than distances of bound steroid substrates. These studies suggest that curcuminoids may cause inhibition of steroid metabolism, especially at higher dosages. Also, the recent popularity of turmeric powder as a dilatory supplement needs further evaluation for the effect of curcuminoids on steroid metabolism. Molecular structure of curcuminoids could be modified to generate better lead compounds with inhibitory effects on CYP17A1 and CYP19A1 for potential drugs against prostate cancer and breast cancer.

## 1. Introduction

Turmeric, the well-known yellow spice and coloring agent, is found in the cuisine of numerous Asian countries. Turmeric is also recognized for its use in Chinese and Ayurvedic medicine and has been tested for anti-microbial activity as far back as 1949 [1, 2]. Turmeric is produced commercially from the dried rhizomes of the plant *Curcuma longa*, which belongs to the ginger family Zingiberaceae. Curcumin (CI-75300, diferuloylmethane, E100, Natural Yellow 3) is the most abundant of the curcuminoids and enhances wound healing, modulates angiogenesis and the immune system; and has anti-inflammatory, anti-oxidant, anti-infective and anti-cancer activities [3]. Since the discovery of curcumin as a bioactive compound, many biological activities have been described [4]. Curcumin has been shown to modulate molecular signaling pathways, such as the aryl hydrocarbon receptor, the induction of Nrf2 or the inhibition of NF-κB, initiating the activation of inflammatory and immunogenic factors. Curcumin can also inhibit angiogenesis and induce apoptosis on cancerous cells [5, 6]. Curcumin is also involved in the increase of insulin in plasma and the decrease of blood glucose in diabetic patients [7, 8]. Curcumin was also found to exhibit a protective effect against natural and chemical toxicities [9]. Structurally, curcumin has two aryl moieties connected by a seven-carbon chain. By varying the motifs from the primary structure, synthetic molecules can be created and a liposomal curcumin preparation can also be prepared for improved stability and bioavailability [10–12].

Anti-cancer activity of curcumin was first described by Kuttan et al. in 1985 [13]. Since then, different research groups have been testing curcumin *in vitro*, and several clinical trials have been carried out to test the biological activities of curcumin preparations [14]. Curcumin molecule itself has poor solubility in water, and therefore, often shows low bioavailability when consumed directly [10, 15]. Different methods have been developed to increase curcumin bioavailability, and in it has been shown that combining curcumin with piperine ((2E,4E)-5-(2H-1,3-Benzodioxol-5-yl)-1-(piperidin-1-yl)penta-2,4-dien-1-one), a component of black pepper obtained from *Piper nigrum*, the bioavailability of curcumin was increased significantly [10, 12, 16].

Most of the hormonal-dependent cancers, such as breast and prostate cancer, are treated by blocking the synthesis of estrogens and androgens [18], by targeting enzymes from the steroidogenesis pathway (Figure 1). The CYP17A1 enzyme (GeneID: 1586, 10q24.32, GRCh38 *chr10:102,830,531-102,837,533*, NCBI: NM_000102.4, NP_000093.1, OMIM: <u>609300</u>) regulates sex steroid biosynthesis in humans through 17α-hydroxylase/17,20 lyase activities and is a target of anti-prostate cancer drug abiraterone [19–21]. Aromatase (CYP19A1) converts androstenedione and testosterone into estrogens and is a target for the treatment for breast cancer [22–24]. The CYP19A1 protein contains 503 amino acids (NP_000094) and is encoded by the *CYP19A1* gene (GeneID:1588, NCBI: NM_000103, 15q21.2, GRCh38 15:51208056-51338597). The cytochrome P450 21-hydroxylase, coded by *CYP21A2* (GeneID: 1589, NCBI: NM_000500.9, NP_000491.4, 6p21.33, GRCh38 chr6:32,038,265-32,041,670, <u>OMIM: 613815</u>) is needed for biosynthesis of mineralocorticoids and glucocorticoids.

**Figure 1:**
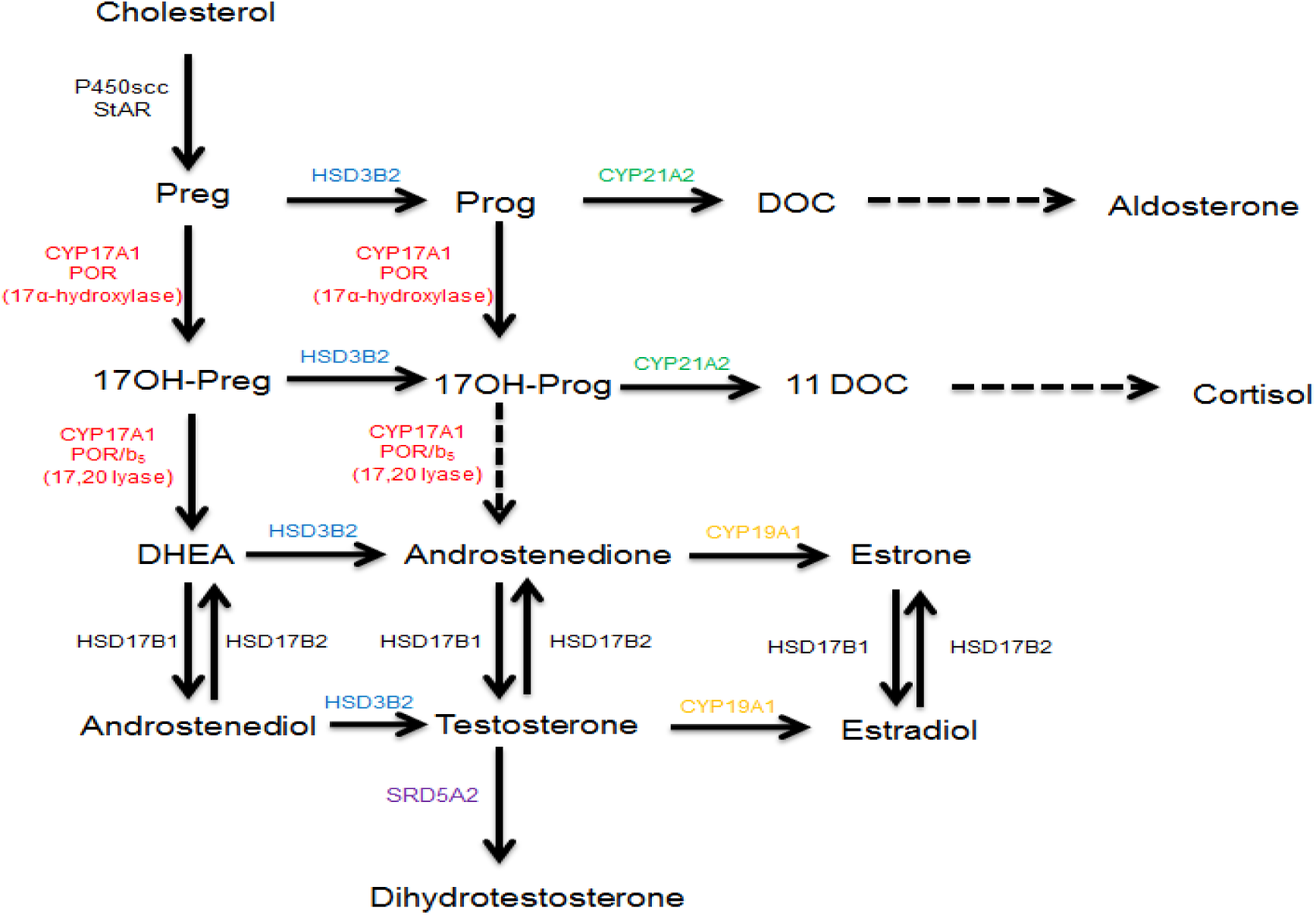
Synthesis of steroid hormones in humans. After entering the mitochondrion, cholesterol is converted into pregnenolone, which is used as a substrate by CYP17A1 in the endoplasmic reticulum to produce sex steroids. First, the cholesterol is converted into pregnenolone by the enzyme CYP11A1 (P450scc) inside the mitochondria. Pregnenolone is converted into progesterone by HSD3B2 and into 17OH-pregnenolone by the 17α-hydroxylase activity of CYP17A1. Progesterone is converted into deoxycorticosterone by CYP21A2 and into 17OH-progesterone by the 17α-hydroxylase activity of CYP17A1. The 17,20 lyase activity of human CYP17A1 converts 17OH-pregnenolone into DHEA but has poor specificity for 17OH-progesterone as a substrate. The 17OH-progesterone is converted into 11-DOC by the 21-hydroxylase activity of CYP21A2. The CYP19A1 (aromatase) converts androgens into estrogens and uses androstenedione as its major substrate. The 5α-reductase, SRD5A2 converts testosterone into dihydrotestosterone. Abbreviations: Preg=pregnenolone, Prog=progesterone, DOC=deoxycorticosterone, 11DOC=11-deoxycortisol, DHEA=dehydroepiandrosterone, HSD3B2=3β-hydroxysteroid dehydrogenase type 2, HSD17B1/2=17β-hydroxysteroid dehydrogenase type 1/2, SRD5A2=5α-reductase type 2.

In the adrenals, CYP21A2 converts progesterone into 11-deoxycorticosterone and 17α-hydroxyprogesterone into 11-deoxycortisol [19]. All these cytochromes P450, the CYP17A1, CYP19A1, and CYP21A2 are membrane-bound proteins and belong to the cytochrome P450 protein superfamily. Cytochromes P450 enzymes are involved in the biotransformation of drugs, xenobiotics, and steroid hormones [25]. There are different types of cytochrome P450 proteins in humans. The cytochrome P450 proteins located inside the mitochondrion metabolize steroids and sterols in partnership with ferredoxin and ferredoxin reductase and are called type 1 cytochrome P450 [26]. The majority of cytochrome P450 proteins in humans (50 out of 57) are located in the smooth endoplasmic reticulum and depend on cytochrome P450 oxidoreductase [27] as their redox partner (type 2 cytochrome P450). The microsomal P450 enzymes metabolize drugs, xenobiotics as well as endogenous substrates, including many steroid hormones like pregnenolone, 17α-hydroxypregnenolone, dehydroepiandrosterone, testosterone, androstenedione [19, 27].

Curcumin, demethoxycurcumin, and bisdemethoxycurcumin are the most abundant components of turmeric; together, these are called curcuminoids (Figure 2). While activities of curcuminoids have been tested against a wide range of metabolic targets, not much is known about the effect of curcuminoids on the steroid metabolism in humans. Considering the widespread use of turmeric powder, which is rich in curcuminoids, we sought to examine the effects of curcuminoids extracted from *C. longa* on the metabolism of steroid hormones. Here we are reporting that curcuminoids may inhibit the biosynthesis of steroid hormones. We tested the effect of curcuminoids at different dosages on the activities of steroid metabolizing enzymes CYP17A1 and CYP21A2 using adrenal carcinoma cell line NCI-H295R and the endoplasmic reticulum from the placental carcinoma cell line JEG3 was used for CYP19A1 activity. Inhibition of CYP17A1 and CYP19A1 activities by curcuminoids indicates similar molecular entities or modified compounds based on core structures of curcuminoids could be explored as potential treatments for prostate cancer by targeting CYP17A1 and for breast cancer by targeting CYP19A1.

**Figure 2:**
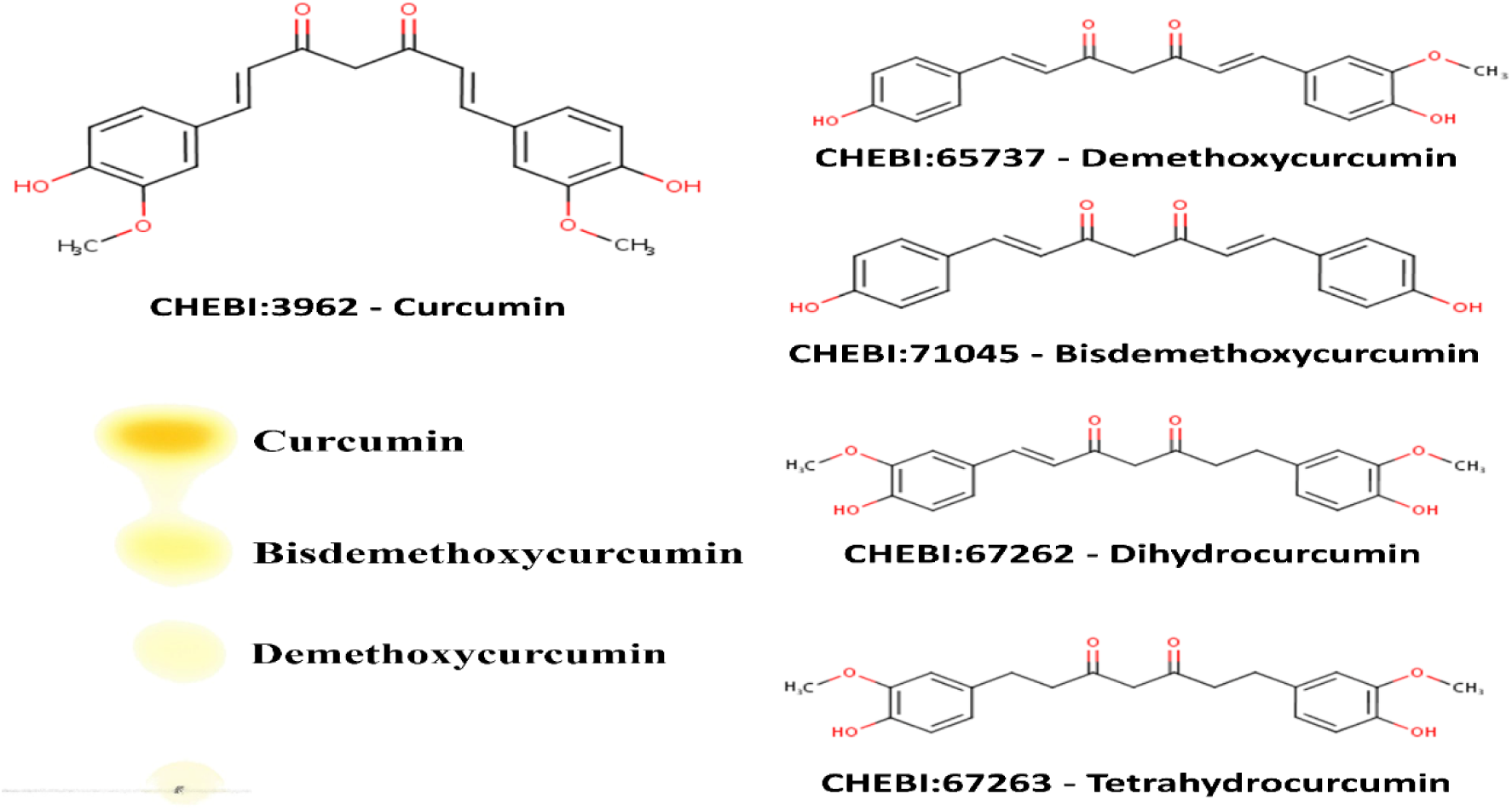
Chemical structure of curcuminoids, and characterization of curcuminoids extracted from turmeric used in this study. The main components of turmeric are curcumin, desmethoxycurcumin, and bis-desmethoxycurcumin. Curcuminoids were obtained from the dried turmeric powder (from *Curcuma longa*) by solvent extraction using ethanol and purified further by selective extraction and crystallization using hexane and isopropanol as described in methods. Curcuminoid composition of our turmeric extract was checked by thin-layer chromatography using chloroform:hexane:methanol (1:1:0.1, v/v/v) as mobile phase. Curcuminoids in our turmeric extract were identified as curcumin (69.7%), bisdemethoxycurcumin (25.8%) and demethoxycurcumin (4.5%) and curcumin was the major component as has been shown in previous publications describing curcuminoid content of *C. longa* [17].

## 2. Results

### 2.1 Isolation of curcuminoids from turmeric powder and characterization

We used dried turmeric powder to obtain the curcuminoid preparation used in our experiments. An initial extraction in ethanol was carried out to dissolve the curcuminoids, and then curcuminoids were separated by filtration. Ethanolic extract was dried under nitrogen and then extracted with hexane to remove bound impurities as curcuminoids are not soluble in hexane. After two rounds of hexane extraction, curcuminoids were crystallized in a 50:50 mix of hexane and propanol and then dried. A stock solution of curcuminoids was made in ethanol. For the determination of the nature of curcuminoids present in our preparation, a thin layer chromatographic separation was carried out as described previously. Our curcuminoid preparation indicated the presence of curcumin as the major curcuminoid (69.7%) and demethoxycurcumin (4.5%) / bisdemethoxycurcumin (25.8%) were present as minor components (Figure 2), which is similar to previous reports describing curcuminoid separation from the powders of *C. longa*.

We performed a UV-Vis spectrum analysis of our curcuminoid preparation and saw an absorption maximum at 427 nm in ethanol (Figure 3A), which was in agreement with values reported previously [28, 29]. Further analysis with fluorescence spectroscopy indicated an emission maximum between 532-538 nm (Figure 3B). Using the known molar extinction coefficient values of purified curcumin (ε=61.864 cm^−1^ mM^−1^), we calculated the total amount of curcuminoids present in our preparation and observed that estimated values were in agreement with the experimentally determined results, indicating high purity of our curcuminoid preparation (Table 1).

**Table 1.** Calculation of curcuminoid concentration based on spectral analysis. Different concentrations of our curcuminoid preparation were analyzed in a SpectraMax M2e spectrophotometer in a volume of 200 µl (pathlength 0.56 cm). The calculated values were in good agreement with the estimated values, indicating a high degree of purity in our curcuminoid preparation.

| Estimated<br>Concentration ($\mu\text{M}$) | $A_{427}$<br>Pathlength 0.56 cm | Observed<br>Concentration ($\mu\text{M}$) |
| --- | --- | --- |
| 1.25 | 0.0477 | 1.4 |
| 2.5 | 0.0915 | 2.6 |
| 5 | 0.1787 | 5.2 |
| 10 | 0.3366 | 9.7 |
| 20 | 0.663 | 19.1 |

**Figure 3.**
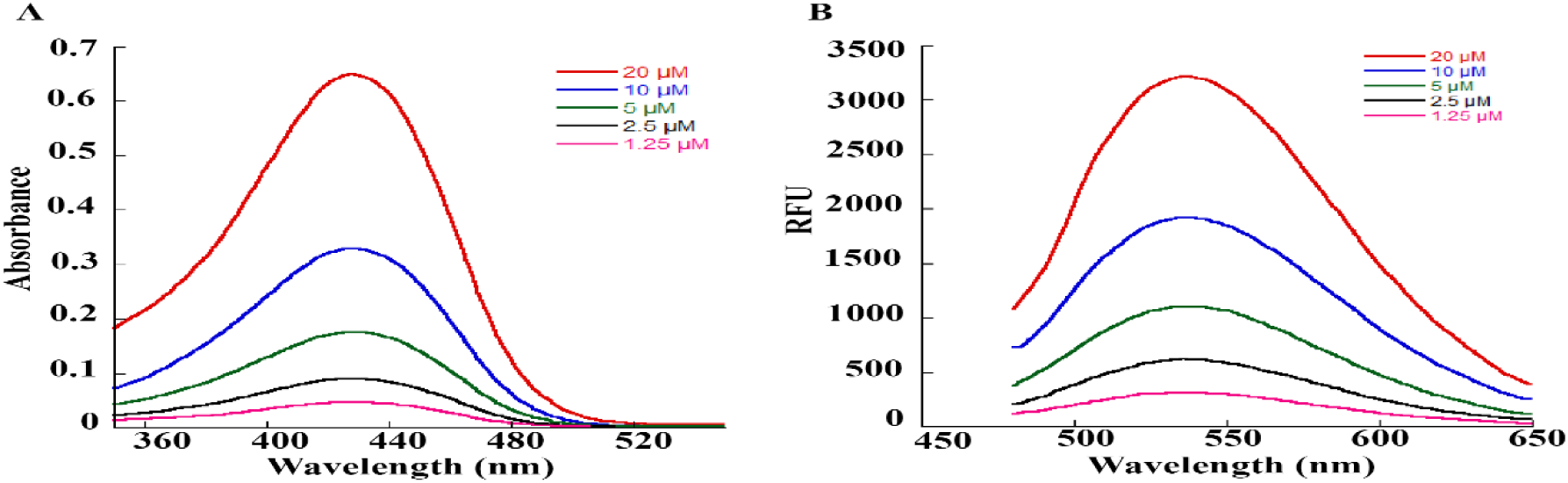
Spectral analysis of curcuminoid preparation used in our studies. A. UV-Vis spectra of curcuminoid preparations. Curcuminoids were dissolved in ethanol and spectra were recorded on a SpectraMax M2e spectrophotometer between 350-550 nm. Different concentrations of the curcuminoid extract (1.25 – 20 µM) were used for recording the spectra. An absorption maximum of 427 nm was observed for our curcuminoid preparation. Based on absorption values obtained from our preparation, an estimation of curcuminoid concentration was performed using a millimolar extinction coefficient of 61.864 cm^−1^ mM^−1^ for pure curcumin (Table 1). B. Fluorescence spectra of increasing concentrations (1.25 – 20 µM) of curcuminoid preparation used in our studies. First, an initial scan of excitation wavelength was performed, then in the emission scans, the excitation wavelength was fixed at 425 nm. A smooth emission maximum between 532-538 nm was observed for our curcuminoid preparation.

### 2.2 Toxicity of curcuminoids on NCI-H295R cells

We first determined the toxicity of the curcuminoids towards the steroid metabolizing NCI-H295R adrenal cancer cell line, which was then used in subsequent experiments (Figure 4). A range of curcuminoid concentrations were tested with NCI-H295R cells to determine the maximum concentration of curcuminoid preparation that was not toxic for the cells. We observed that at 50 µg/ml or higher concentrations of curcuminoid preparation, only 25% of the NCI-H295R cells survived, while at 25 µg/ml or lower concentrations of curcuminoids, most of the cells were found to be viable. We did not observe cell death between 0.78-12.5 µg/ml of curcuminoid preparation, and a standard concentration of 10 µg/ml curcuminoid preparation was used in further experiments. Similarly, the toxicity of curcuminoid preparation was also tested for HEK-293 cells, and no significant toxicity was observed below 6.25 µg/ml or higher concentrations (Supplementary Figure 1). These results are in agreement with the previously reported toxicity values for curcumin in NIH3T3, H9C2, and HepG2 cells [30]. Human trials using up to 8000 mg of curcumin found no evidence of toxicity [31].

**Figure 4:**
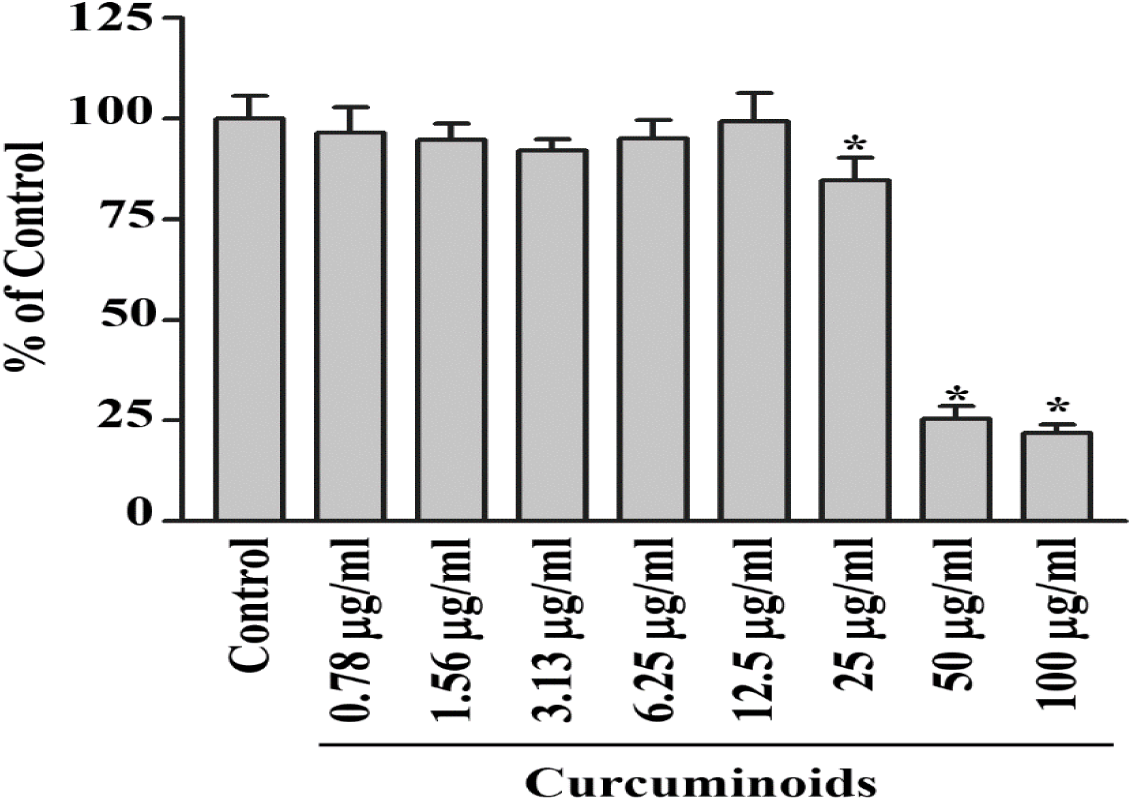
Measurement of cytotoxicity of curcuminoid preparation. Effect of curcuminoids on cytotoxicity and viability of human adrenal NCI-H295R cells was determined using a range of curcuminoid concentrations between 0.78-100 µg/ml over 24 h as described in methods. The NCI-H295R cells were grown overnight and then treated with varying concentrations of curcuminoids for 24 h. After the incubation with curcuminoids, culture medium was removed, and cell viability was determined by MTT reduction assay. No significant effect on the viability of NCI-H295R cells was observed at 25 µg/ml or lower doses of curcuminoids. In subsequent experiments, the concentration of curcuminoids was kept below 10 µg/ml. Data are presented as the mean and standard deviation of 3 independent replicates.

### 2.2 Bioactivity of curcuminoids on steroid biosynthesis

We tested the bioactivity of our curcuminoid preparation of steroid production by human adrenal NCI-H295R cells [32]. The NCI-H295 cells have been shown to express the enzymes involved in the biosynthesis of steroids in human adrenals and therefore, the NCI-H295R cells have been an excellent model to study the molecular mechanisms of steroid regulation and human adrenal steroidogenesis [33]. To test the effects of curcuminoids on adrenal steroid production, we treated the NCI-H295R cells with 10 µg/ml of curcuminoid preparation and used pregnenolone as a substrate. We compared the effect of curcuminoids on steroid biosynthesis in NCI-H295R cells with abiraterone, a known inhibitor of CYP17A1 and CYP21A2 activities. The curcuminoids, as well as abiraterone, blocked the production of 17α-hydroxypregnenolone and DHEA, indicating inhibition of CYP17A1 activities (Figure 5).

**Figure 5.**
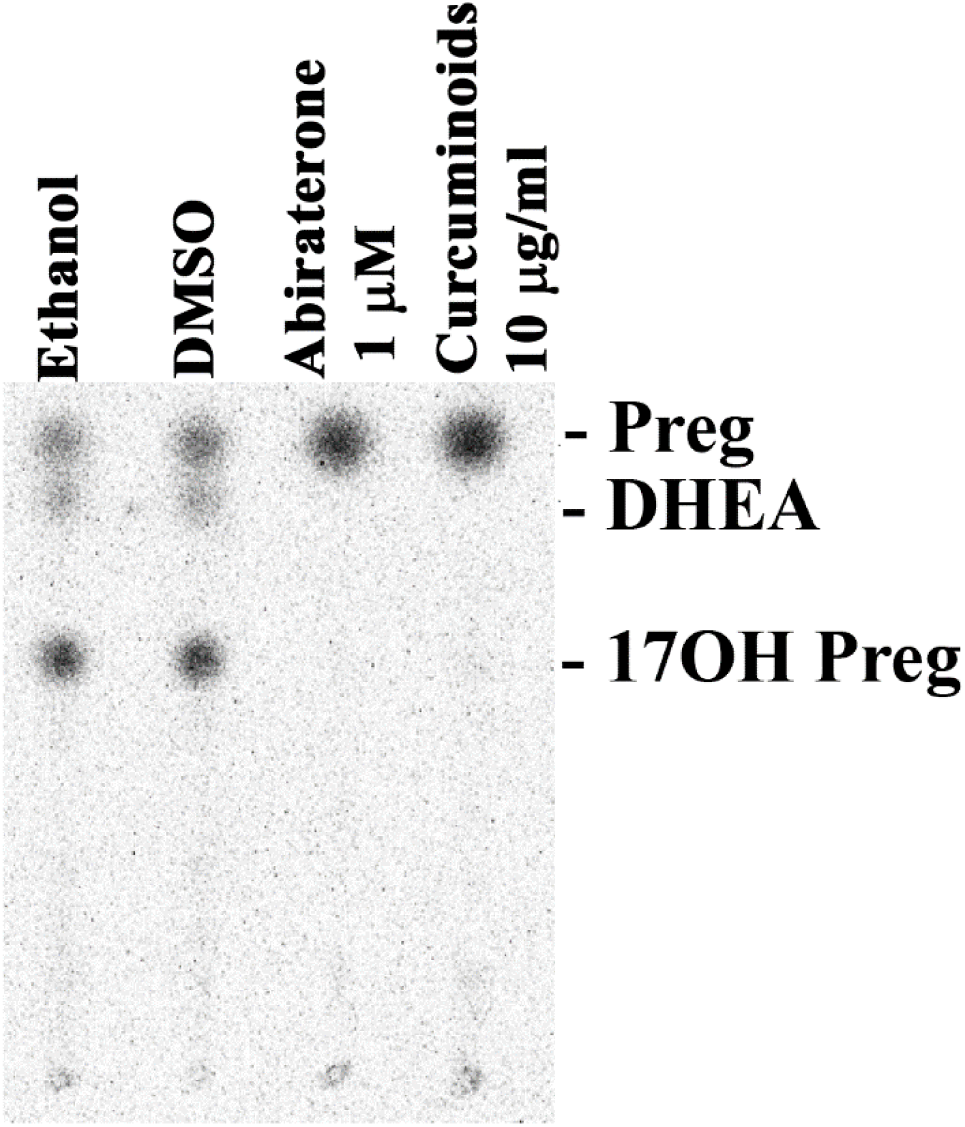
Effect of curcuminoids on steroid production. A representative thin layer chromatogram is shown illustrating the effect on steroid production by the NCI-H295R cells from the curcuminoids extracted from turmeric. Human Adrenal NCI-H295R cells were incubated with tritium labeled pregnenolone and then treated with curcuminoids and abiraterone, a known inhibitor of CYP17A1. After incubations, steroids were extracted by organic solvents, dried under nitrogen and separated by TLC. Radioactivity contained in steroids was visualized using autoradiography on a phosphorimager and quantitated by densiometric analysis. Curcuminoids inhibited the formation of 17OH-pregnenolone and DHEA in human adrenal NCI-H295R cells. A block of DHEA production indicates that curcuminoids caused inhibition of both the 17α-hydroxylase as well as 17,20 lyase activities of CYP17A1. Ethanol and DMSO were used as controls at 0.1% of the total reaction volume.

### 2.3 Bioactivity of curcuminoids on CYP17A1

After we saw the preliminary results indicating a block of DHEA production by curcuminoids that were similar to inhibition caused by abiraterone, we did further experiments to check the details of steroid biosynthesis inhibition by curcuminoids. We tested the effect of different concentrations of curcuminoids on the 17α-hydroxylase and 17,20 lyase activities of CYP17A1 (Figure 6A). We observed inhibitory effects of curcuminoids on both the 17α-hydroxylase as well as 17,20 lyase activities of CYP17A1 in a dose-dependent manner (Figure 6B and 6C). Abiraterone, a known inhibitor for CYP17A1, which was used as a control, also inhibited the CYP17A1 activities. Overall, inhibition by curcuminoids of the CYP17A1 17, 20 lyase activity seemed stronger compared to the inhibition of the 17α-hydroxylase activity (Figure 6C). A stronger inhibition of 17,20 lyase activity may be due to direct competition with 17OH-pregnenolone or blocking the interaction of CYP17A1 with cytochrome b_5_ [34].

**Figure 6:**
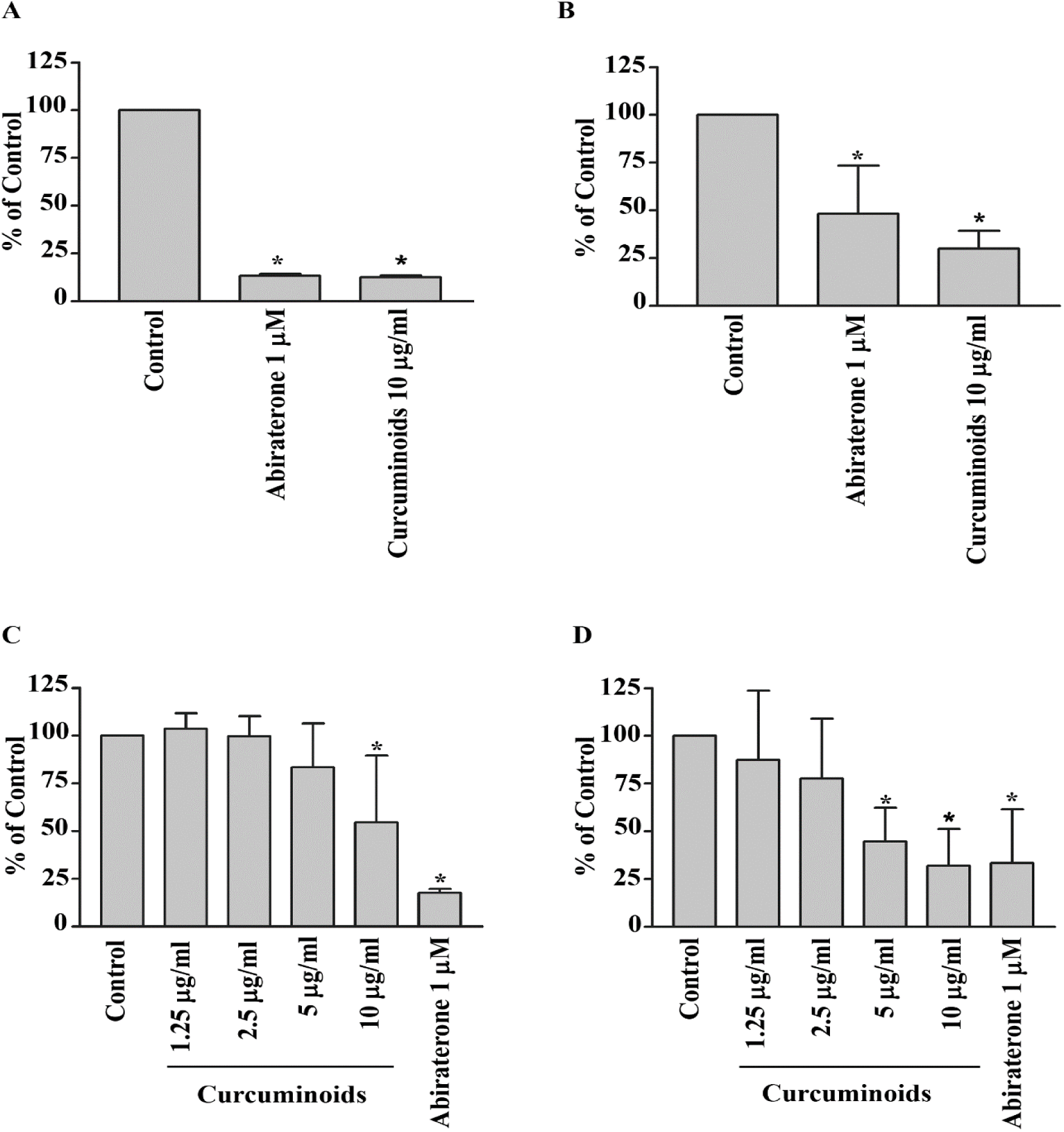
Inhibitory effects of curcuminoids on CYP17A1 activities. Abiraterone, a known inhibitor for CYP17A1, was used as a control. Human adrenal NCI-H295R cells were treated with different concentrations of the curcuminoid preparation extracted from turmeric. The CYP17A1 converts pregnenolone into 17α-hydroxypregnenolone by its 17α-hydroxylase activity, and 17α-hydroxypregnenolone is converted into DHEA by the 17,20 lyase activity of CYP17A1. Each experiment was done in triplicate. The final volume of ethanol in experiments was 0.1%, and trilostane (at 1µM concentration) was used to block the activity of HSD3B2, which converts pregnenolone into progesterone. First a pilot experiment was performed to test the effect of curcuminoids (A and B), and then several different concentrations of curcuminoids were used to determine the inhibition of CYP17A1 activities by curcuminoids (C and D). A. Inhibition of CYP17A1 17α-hydroxylase activity at a curcuminoid concentration of 10 µg/ml, B. Inhibition of CYP17A1 17,20 lyase activity at a curcuminoid concentration of 10 µg/ml. C. Differential inhibition of curcuminoids on CYP17A1 17α-hydroxylase activity. D. Differential inhibition of curcuminoids on CYP17A1 17, 20 lyase activity. Data are presented as the mean and standard deviation (error bars) of 3 independent replicates with p values < 0.05 (*) considered significant. Student t-test was used to calculate the differences between samples.

### 2.4 Bioactivity of curcuminoids on CYP21A2

We used 17-hydroxyprogesterone as a substrate for CYP21A2 activity and monitored the production of 11-deoxycortisol to quantify the effect of curcuminoids. Human adrenal NCI-H295R cells do not use 17-hydroxyprogesterone as a substrate of CYP17A1 and therefore, are a good model for assay of CYP21A2 activities using 17-hydroxyprogesterone. We tested the bioactivity of curcuminoids at 5 µg/ml and 10 µg/ml concentrations for effect on CYP21A2 activity (Figure 7). At 10 µg/ml concentration of curcuminoids the activity of CYP21A2 was only moderately reduced compared to control, and was less than the effects observed for inhibition of CYP171 and CYP19A1 activities. The anti-prostate-cancer drug abiraterone also inhibited the CYP21A2 activity as we have shown previously [20, 21].

**Figure 7:**
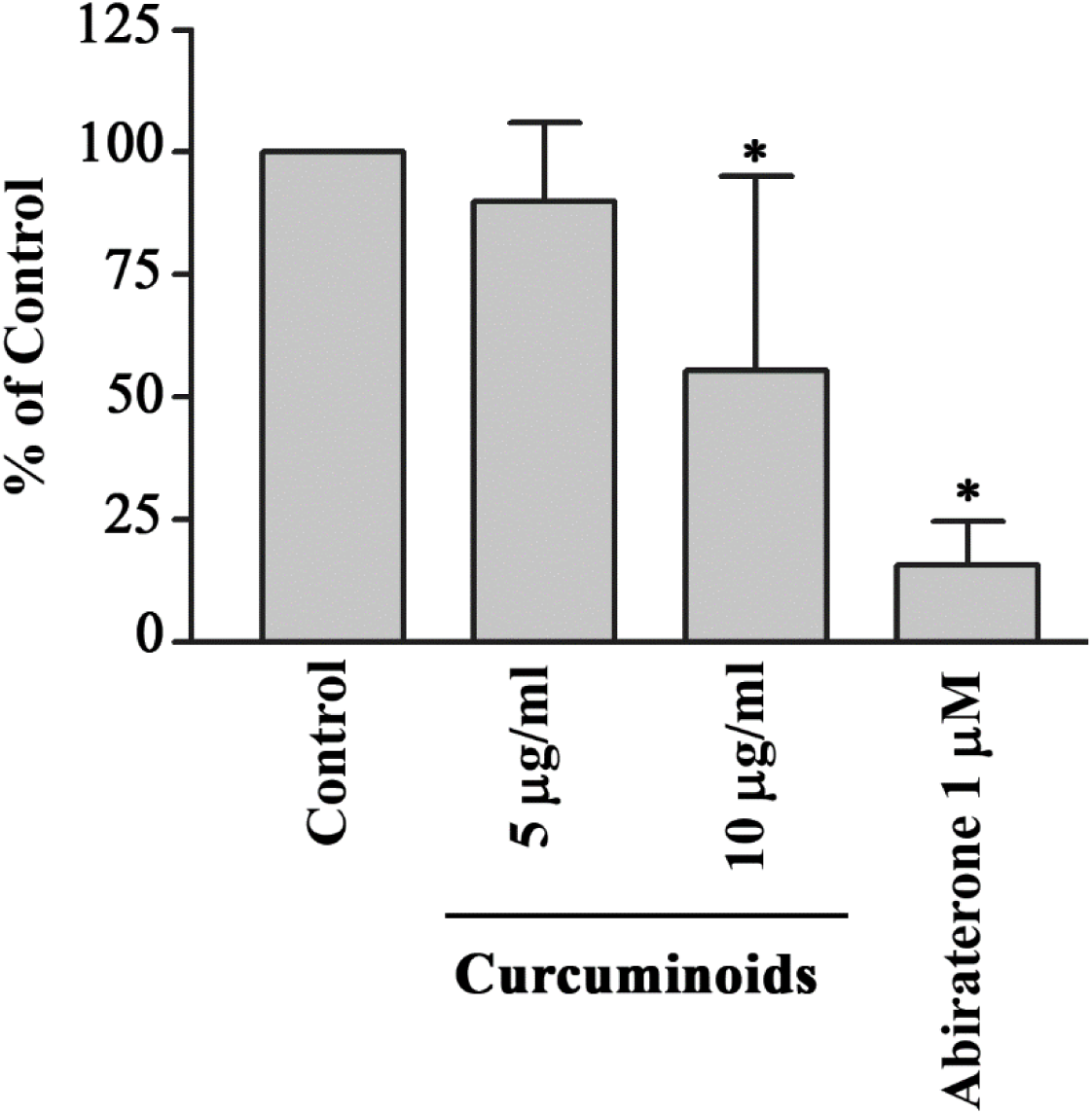
Effect of curcuminoids on CYP21A2 activity. Bioactivity of curcuminoids on CYP21A2 was measured at two different concentrations. Human adrenal NCI-H295R cells were used as a source of CYP21A2 activity. In the human adrenals, CYP17A1 does not utilize 17α-hydroxyprogesterone as a substrate, and therefore, for the assay of CYP21A2 activity, 17α-hydroxyprogesterone was used as substrate and reaction was monitored by measuring the production of 11-deoxycortisol. NCI-H295R cells were seeded overnight and then incubated with curcuminoids. After the incubations, [^3^H]-17α-OH progesterone was added to the medium as the substrate. Following the reactions, steroids were extracted by organic solvents, dried under nitrogen and separated by TLC. Production of 11-deoxycortisol from 17α-OH progesterone was measured by autoradiographic analysis of steroids on TLC plates as described in methods. The activity of CYP21A2 was moderately reduced at 10 µg/ml of curcuminoids. Ethanol concentration was 0.1% in the control reaction and abiraterone, a known inhibitor of CYP21A2 was used as a positive control. Data shown here are from the mean and standard deviation (error bars) of three independent experiments.

### 2.5 Bioactivity of curcuminoids on CYP19A1

To test the effect of curcuminoids on the aromatase activity of CYP19A1, we used androstenedione as substrate and monitored its metabolism by quantifying the release of tritiated water from radiolabeled androstenedione. A preparation of endoplasmic reticulum obtained from the placental JEG3 cells was used for the assay of aromatase enzyme activity. A dose-response effect showing the inhibition of aromatase with the increasing amounts of curcuminoids was seen from 0.78 to 100 µg/ml concentration of curcuminoids (Figure 8). A known inhibitor of CYP19A1, anastrozole was used as a positive control at 100 nM concentration and showed inhibition of CYP19A1 activity as expected.

**Figure 8:**
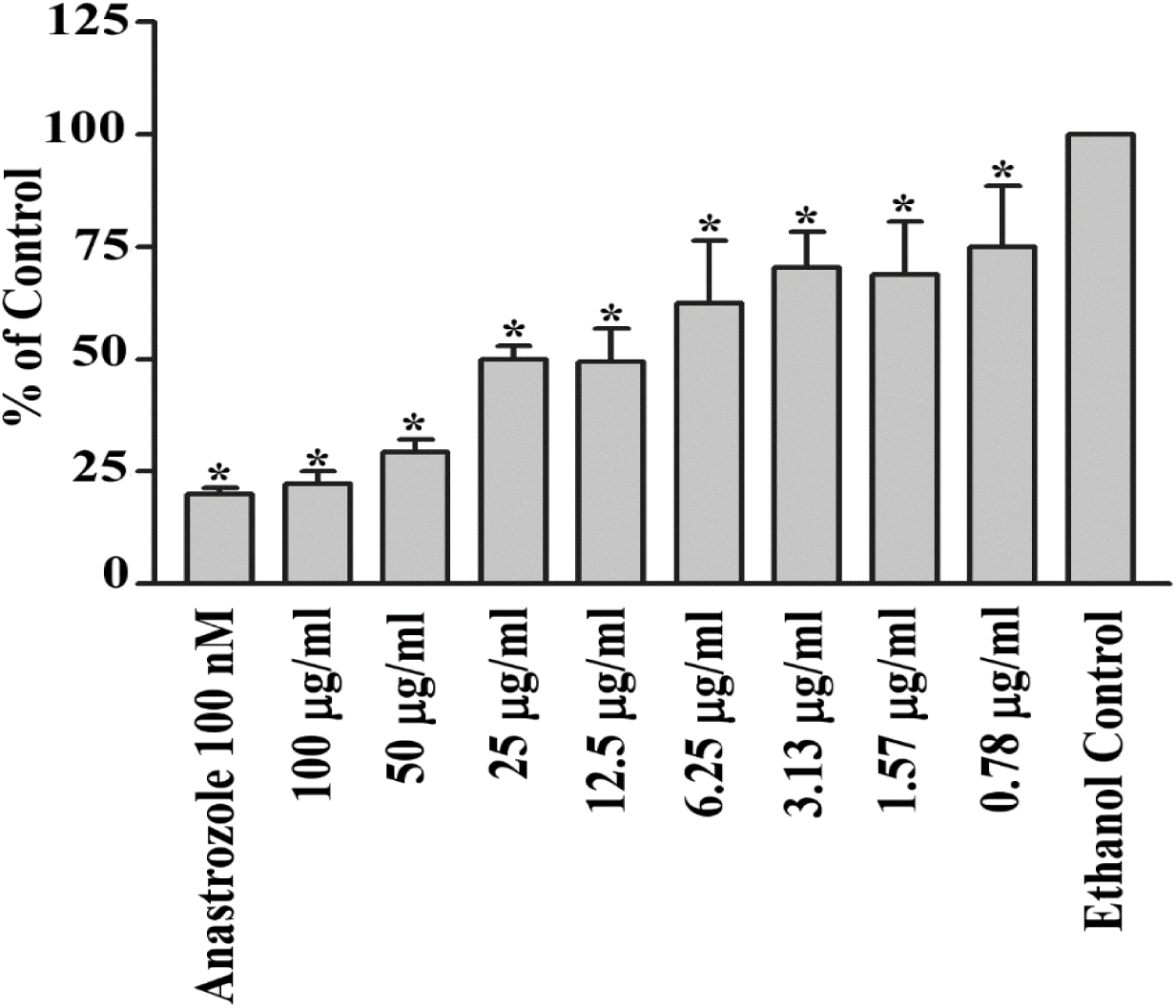
Effect of curcuminoids on CYP19A1 activity. A dose-response profile is shown indicating the inhibition of aromatase by curcuminoids. A preparation of endoplasmic reticulum obtained from JEG3 placental cells was used as a source of aromatase activity, and tritium-labeled androstenedione was used as a substrate. The CYP19A1 reaction was monitored by calculating the amount of tritiated water released by CYP19A1 during the aromatization of androstenedione. A known inhibitor of CYP19A1, anastrozole (100 nM) was used as a positive control. Curcuminoids showed a small effect on CYP19A1 activity 6.25 µg/ml and higher inhibition was observed at 12.5, 25, 50 and 100 µg/ml concentrations of curcuminoids, indicating natural curcuminoids present in *C.longa* are not potent inhibitors of aromatase activity. Data are presented as the mean and standard deviation of 3 independent replicates. Student t-test was used to calculate the differences between samples (* p values < 0.05).

### 2.6 Computational docking of curcumin into the human CYP17A1, CYP21A2, and CYP19A1 crystal structures

After observing the inhibitory effects of curcuminoids on CYP17A1, CYP19A1, and CYP21A2, we performed computational docking of curcumin into the protein structures of these cytochromes P450 to understand the molecular nature of inhibition. Molecular structure of curcumin resembles steroid substrates of cytochromes P450 studied in this report, and therefore, we wanted to check whether curcumin fits into the active site of these steroid metabolizing enzymes. Curcumin was docked into the crystal structures of human steroid metabolizing cytochrome P450 CYP17A1, CYP19A1, and CYP21A2 using the docking program Autodock VINA (Figure 9). Superimposition of P450 structures with either their substrates or the curcumin docked into the active site revealed similar binding poses (Figure 9). We observed a close binding pattern from the docking of curcumin into the active sites of CYP17A1 and CYP19A1 crystal structures (Heme iron to curcumin distances < 2.5 Å) (Figure 9). The binding pose of curcumin into CYP21A2 structure was also like its binding into the CYP17A1 and CYP19A1 structures.

**Figure 9:**
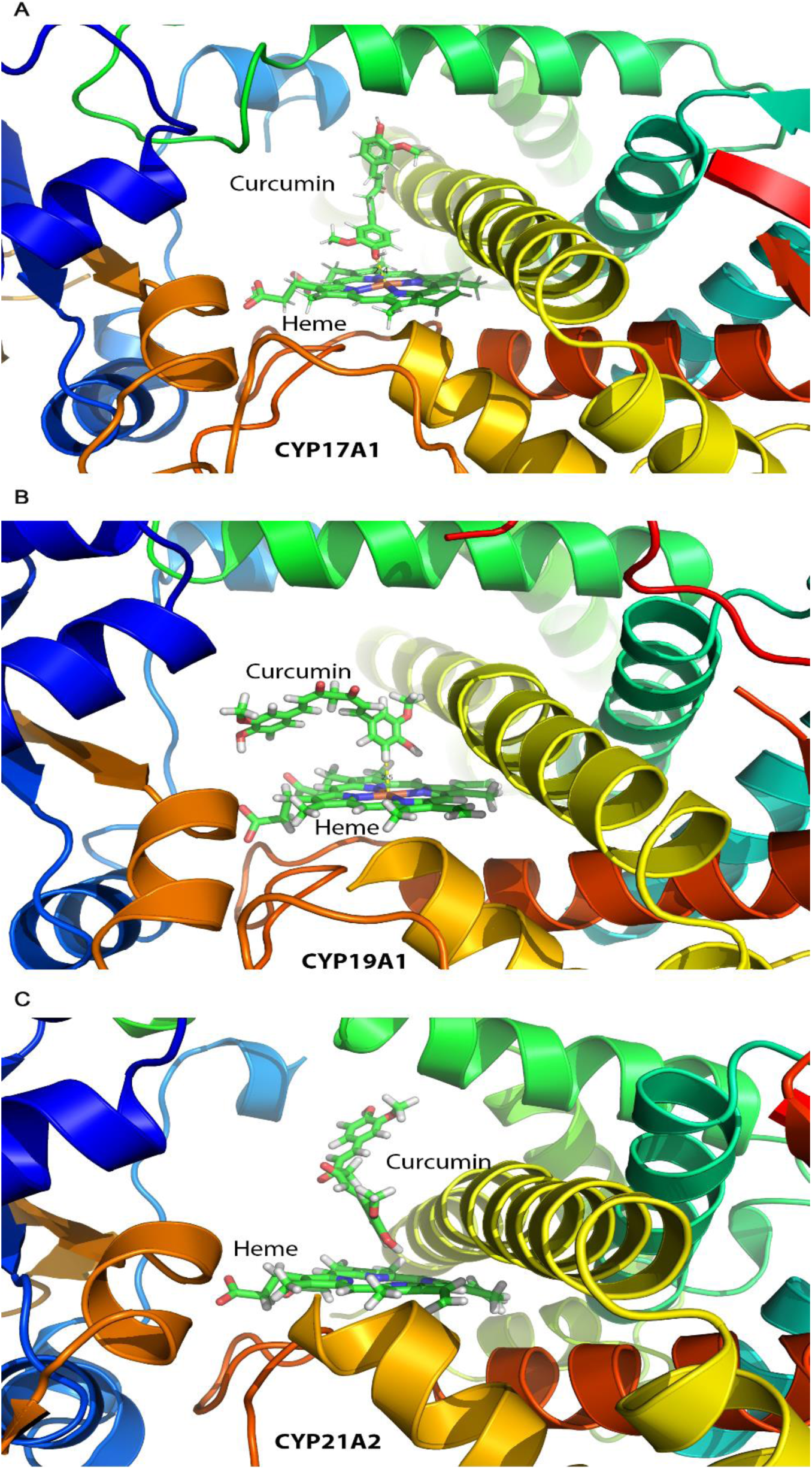
Docking of curcumin into the protein structures of CYP17A1, CYP19A1, and CYP21A2. Published crystal structures of CYP17A1, CYP19A1, and CYP21A2 were used for docking of curcumin by software AUTODOCK-VINA as described in materials and methods. Bound steroid ligands were removed before docking of curcumin into the active site of P450s. The poses in which curcumin docks are similar to steroid substrates of all three P450s, with closer fitting in case of CYP17A1 and CYP19A1 as compared to CYP21A2 (distance from heme 2.5 Å for CYP17A1/CYP19A1 versus 3.4 Å for CYP21A2).

A comparison of the CYP17A1 and CYP19A1 crystal structures in complex with curcumin and docked curcumin into the crystal structure of CYP21A2 revealed similar binding conformations but the distance of the curcumin to the central heme iron of CYP21A2 was longer (3.4 Å *versus* 2.4 Å) (Figure 9). Binding of curcumin with CYP17A1, CYP19A1 and CYP21A2 shares many similarities with the binding of steroid substrates into the active sites of these cytochrome P450 proteins with many similar active site residues involved in binding. These results indicate that curcumin may act as an active site inhibitor and occupy the same space in cytochrome P450 active sites as used by their steroid substrates.

## 3. Discussion

All steroid hormones are produced from cholesterol [35]. The abnormal production of steroid hormones, for example, the hypercortisolemia seen in the Cushing’s syndrome, is a life-threatening condition [36, 37]. The hyperandrogenism is not life-threatening by itself but creates many severe complications during all phases of life. Androgens regulate sexual differentiation in both the female as well as male and disruption of androgen biosynthesis by mutations in steroid metabolizing enzymes, or their redox partners, cause metabolic disorders [38]. Non-tumoral cases of hyperandrogenism include the polycystic ovary syndrome (most common metabolic disorder in females), congenital adrenal hyperplasia caused by to 21-hydroxylase deficiency and Cushing syndrome [39, 40]. Overproduction of cortisol and androgens could be therapeutically controlled by inhibitors that block specific steps in steroid biosynthesis.

CYP17A1 catalyzes multiple reactions in the steroid biosynthesis pathway [41–43]. Main activities of CYP17A1 include the 17α-hydroxylase activity needed for the biosynthesis of 17OH-pregnenolone (17OH-PREG) and 17OH-progesterone (17OH-PROG), the precursors of cortisol. The CYP17A1 provides qualitative regulation of steroid production in humans by its 17,20 lyase activity that produces dehydroepiandrosterone (DHEA), the precursor of sex steroids. The two distinct enzyme activities of the CYP17A1 dictate the nature of steroids synthesized in different types of cells [19, 34]. Overproduction of androgens by the activation of CYP17A1-17,20 lyase activity has been linked to polycystic ovary syndrome. CYP17A1 is also a metabolic target for chemotherapy of castration-resistant prostate cancer [20, 44].

Novel compounds that are safe and non-toxic are needed to target CYP17A1 and CYP19A1 activities to treat metabolic disorders resulting from excess production of androgens or estrogens. Here we have probed the potential of curcuminoids as possible test candidates for synthesizing novel chemicals to target CYP17A1 and CYP21A2. We found a dose-dependent inhibition of both the CYP17A1 and CYP19A1 by curcuminoids. In our current study, curcuminoids caused a reduction of CYP21A2 activity. A reduction in CYP21A2 activity was observed at 10 µg/ml concentration of curcuminoids. Based on these results, we can conclude that curcuminoids may cause an intricate pattern of changes in steroid metabolites due to small to moderate inhibition of CYP19A1 and CYP21A2 activities in addition to significant inhibition of 17α-hydroxylase as well as 17,20 lyase activity of CYP17A1.

Curcumin has been tested on the five drug-metabolizing enzymes from the cytochrome P450 family, *in vivo* [45] and *in vitro* [46, 47], showing that CYP1A2, CYP2B6, CYP2C9, CYP2D6, and CYP3A4 can be inhibited by curcuminoids in a dose-dependent matter. Our studies provide an analysis of the inhibitory effects of curcuminoids on CYP17A1 and CYP19A1 activities. These results indicate that the steroid production in people with high amounts of Curcuma consumption may be affected not only by the inhibition of CYP17A1 activities but also by the inhibition **of** CYP19A1 and to a smaller extent, the inhibition **of** CYP21A2 enzyme activity. Also, use of curcuminoids as a common over the counter health supplement requires further caution as inhibition of both the CYP17A1 as well as the CYP19A1 activities, may potentially result in complications of steroid metabolism in both males and females. Inhibitors of steroidogenesis act on multiple steps in steroid biosynthesis. Many drugs have been approved for use as steroid biosynthesis inhibitors (Ketoconazole, Metyrapone, Etomidate, Mitotane). All these drugs inhibit CYP17A1, CYP11A1, CYP11B1, and CYP19A1 [48]. Also, many **other** inhibitors which can block androgen biosynthesis in androgen-dependent prostate cancers are being studied [21, 49–51].

Computational docking from our experiments showed that curcumin could bind into the active sites of steroid metabolizing P450s. The inhibition of CYP17A1 and CYP19A1 by curcuminoids provides a template for structure modification to produce effective and safe compounds that can target prostate cancer as well as breast cancer. Only a minor effect on CYP21A2 activity by curcuminoids compared to inhibition of CYP17A1 (50% inhibition of 17,20 lyase activity at 5 µg/ml compared to no significant effect on CYP21A2 activity at the same concentration) was observed. This suggests that compounds based on curcuminoids may be better and safer inhibitors for use in hyperandrogenic states like polycystic ovary syndrome, especially in children and young adults, and avoid the toxic effects of abiraterone which severely inhibits CYP21A2 activity [20, 21].

## 4. Materials and Methods

### 4.1 Materials

Radiolabeled [^3^H]-pregnenolone [7(N)-^3^H]-pregnenolone (12.6 Ci/mmol) and [^3^H]-Androstenedione: Andros-4-ene-3, 17-dione, [1β-^3^H(N)]-, 250 µCi (9.25 MBq) were purchased from PerkinElmer (Waltham, MA, USA). [^3^H]-17α-OH progesterone [1, 2, 6, 7-3H] (60-120 Ci/mmol 2.22-4.44 TBq/mmol) was obtained from American Radiolabeled Chemicals Inc. (St. Louis, MO, USA). Silica gel-coated aluminum backed TLC plates were purchased from Macherey-Nagel (Oensingen, Switzerland). The tritium-screens used for the autoradiography were purchased from Fujifilm (Dielsdorf, Switzerland). Turmeric extract capsules (from *Curcuma longa)* were obtained from the Finest Natural (Item: 943.17, Deerfield, IL, USA). Trilostane was extracted in absolute ethanol (EtOH) from tablets commercially available as Modrenal® (Bioenvision, NY, USA). Abiraterone was purchased from Selleckchem (Houston, TX, USA). Anastrozole was obtained from AstraZeneca (Cambridge, UK).

### 4.2 Cell lines and culture media

Human placental JEG3 cells were purchased from American Type Culture Collection (ATCC) (ATCC: HTB-36™) and cultured in MEM with Earle’s salts (GIBCO) supplemented with 10% Fetal Bovine Serum, 1% L-Glutamine (200 mM GIBCO), 1% penicillin (100 U/ml; GIBCO) and streptomycin (100 µg/ml; GIBCO). Human adrenocortical NCI-H295R (NCI-H295R) cells were purchased from ATCC (ATCC: CRL-2128) and grown in DMEM/Ham’s F-12 medium containing L-glutamine and 15 mM HEPES (GIBCO, Paisley, UK) supplemented with 5% NU-I serum (Becton Dickinson, Franklin Lakes, NJ USA), 0.1% insulin, transferrin, selenium (100 U/ml; GIBCO), 1% penicillin (100 U/ml; GIBCO) and streptomycin (100 μg/ml; GIBCO). HEK293 cells were grown in DMEM GlutaMAX^TM^ medium supplemented with 10% FBS, 1% antibiotic mix (100x) and 1 mM sodium pyruvate.

### 4.3 Curcuminoid extraction and analysis

To obtain a crude turmeric extract, we mixed 20 g of turmeric in 200 ml of ethanol and kept it overnight. The ethanol extract was filtered and dried under nitrogen. Pure curcuminoids were obtained by recrystallization using hexane and propanol as described previously. In brief, dried ethanol extract powder was dissolved in hexane and centrifuged at 2000 x g for 15 minutes, and the supernatant was discarded. The pellet from hexane extraction was dissolved in a propanol/hexane mixture (50% propanol + 50% hexane). The solution was mixed for 90 min and then stored at 4 °C overnight for crystallization. The next day, soluble fraction was removed, and the crystallized curcuminoids were dried under nitrogen. A stock solution of curcuminoid stock was made by mixing 10 mg of the crystallized curcuminoid powder in 1 ml of 100% ethanol. The purity of the curcuminoid extract was tested by thin-layer chromatography [52] (Figure 2). Curcuminoid extract was further analyzed by UV-Vis and fluorescence spectroscopy to compare the spectral properties of our preparation with previous extraction procedures.

The Absorption spectra of curcumin in ethanol were collected using a microplate reader (Spectramax M2e, Molecular Devices, Sunnyvale, CA). In brief, different concentrations of curcumin (1.25–20 μM) in ethanol were scanned within the wavelength range of 350–550 nm at 1 nm interval in a total volume of 200 μl. All UV–Vis measurements were performed at 25 °C and corrected for the solvent medium (ethanol). Similarly, the fluorescence spectra of varying concentrations of curcuminoids were also recorded at 25 °C on a microplate spectrofluorometer system (Spectramax M2e, Molecular Devices, Sunnyvale, CA). Ethanol solutions of curcuminoids at 1.25, 2.5, 5, 10 and 20 μM concentration in a volume of 100 μl were scanned at 2 nm interval. The emission spectra were measured with an excitation wavelength set at 425 nm, and emission was scanned between 480-650 nm.

### 4.4 Cell viability assay using MTT

Human adrenal NCI-H295R cells were seeded in 96-well culture plates at a density of 3×10^5^ cells per well and grown overnight at 37°C under 5% CO_2_, and 90% humidity. After 24 h, the medium was changed, and serial dilutions of curcuminoids were added to the medium and incubation of cells was continued for another 24 h. After the incubation of cells with curcuminoids, 20 µl of MTT reagent (5 mg/ml in PBS) was added into each well, and the incubation was continued for another 4 h. After the incubation with MTT reagent, culture medium was removed and 200 µl of DMSO was added in each well, and the plate was incubated for 20 min in the dark. The absorbance in individual wells was then measured at 570 nm, and calculations of cell viability were performed based on residual MTT reduction activity of cells compared to controls without curcuminoids. As another control, the toxicity of curcuminoids on human HEK-293 cells was also tested under similar conditions using the methods described for NCI-H295R cells.

### 4.5 Assay of CYP17A1 and CYP21A2 activities

Human adrenal NCI-H295R cells were seeded in 6 well tissue culture plates at a density of 1×10^6^ cells per well. After overnight incubation, medium was changed, and different concentrations of curcuminoids were added to incumation medium and incubation was continued for 24 h. A known inhibitor of CYP17A1 and CYP21A2, Abiraterone (1 µM), was used as a control for both assays. NCI-H295R cells were treated with trilostane (1 µM), an inhibitor of HSD3B2, 90 min before the addition of radiolabeled substrates. For CYP17A1 activity assays, ~100000 cpm/well of [^3^H]-pregnenolone was added to each well. For the determination of CYP21A2 activity, [^3^H]-17α-OH progesterone (~50000 cpm/well) was used as a substrate. After incubations with radiolabeled substrates, medium from each well was collected, and steroids were extracted in ethyl acetate:isooctane (1:1). Extracted steroids were concentrated under nitrogen and dissolved in 20 µl of trichloromethane for separation by thin-layer chromatography on silica gel TLC plates (Macherey-Nagel, Oensingen, Switzerland). Radiolabeled steroids were quantified by autoradiography on a Fuji FLA-7000 PhosphorImager (Fujifilm, Dielsdorf, Switzerland). Quantification of steroids was done using MultiGauge software (Fujifilm, Dielsdorf, Switzerland).

### 4.6 Preparation of microsomes from JEG3 cells

JEG3 cells were collected near confluency and washed with cold PBS. The cell suspension was then centrifuged at 1500 x g for 5 min to pellet the cells. Afterward, the cell pellet was suspended in 100 mM Na_3_PO_4_ (pH 7.4) containing 150 mM KCl, and the cells were lysed by sonication. Unbroken cells and mitochondria were pelleted by centrifugation at 14000 x g for 15 min at 4°C. Microsomes containing endoplasmic reticulum were collected by ultracentrifugation at 100000 x g for 90 min at 4°C and resuspended in 50 mM K_3_PO_4_ (pH 7.4) containing 20% glycerol [34]. The protein content of microsomes was measured by the BioRad Protein Assay Kit (BioRad, Hercules, CA, USA).

### 4.7 Assay of CYP19A1 activity

The aromatase activity of CYP19A1 was measured by calculating the release of tritiated water from the substrate during aromatization. Assays of aromatase activity were carried out using 40 µg of microsomes from JEG3 cells in 100 mM potassium phosphate buffer (pH 7.4) containing 100 mM NaCl in a final volume of 200 µl. Different concentrations of curcuminoids were added to reaction mixtures, and ethanol was used as control. A known inhibitor of CYP19A1, anastrozole was included in some reactions as a positive control. Assays were performed using 50 nM androstenedione as a substrate and contained ~20000 cpm of [^3^H]-Androstenedione as a radioactive tracer. The aromatase reaction was initiated by addition of NADPH to 1 mM final concentration and incubations were done at 37°C with constant shaking for 1h. Afterward, 800 µl of a solution containing 5% charcoal with 0.5% dextran was added to each reaction mixture and mixed by vortexing followed by centrifugation at 15000 x g for 10 min. From each reaction, 500 µl aliquots were taken for the measurement of radioactivity by scintillation counting using the Rotiszint Universal Cocktail (Carl Roth GmbH, Karlsruhe, Germany), and aromatase activity was calculated as described previously [53, 54].

### 4.8 Docking of curcumin into CYP17A1, CYP19A1, and CYP21A2 protein structures

The published 3D structures of human CYP17A1, CYP19A1, and CYP21A2 were obtained from the PDB database (www.rcsb.org). We made in-silico calculations and structure analysis with YASARA [55]. For docking experiments, X-ray crystal structures of human of CYP17A1 (PDB# 3RUK), CYP19A1 (PDB# 3EQM) and CYP21A2 (PDB # 4Y8W), were used [56–58]. Missing hydrogen atoms into the structures were added with YASARA [55] which was also used for all other calculations. The resulting minimum energy structures were used for AutoDock VINA [59] to perform docking experiments with curcumin (orthorhombic docking was grid established around the central heme molecule). The final poses of curcumin were selected based on their docking location and scores, and resemblance to the co-crystallized ligands in the P450 structures. Structure models were drawn with Pymol (www.pymol.org) and rendered as ray-traced images with POVRAY (www.povray.org).

### 4.9 Statistical analysis

For the statistical analysis, Microsoft Excel and GraphPad Prism (Graph Pad Software, Inc. San Diego, CA, USA) were used. Data are presented as the mean and standard deviation of 3 independent replicates. Student t-test was used to calculate the differences between experiments. Significance cutoff for all experiments was set at p < 0.05.

